# Altered metabolic health during pregnancy in mice with lean polycystic ovary syndrome-like traits from high prenatal AMH

**DOI:** 10.64898/2026.02.26.708393

**Authors:** Emma J. Houston, Emily Jewett, Faria Athar, Nicole M. Templeman

**Affiliations:** Department of Biology; University of Victoria; Victoria, British Columbia, Canada; V8P 5C2

## Abstract

Polycystic ovary syndrome (PCOS) is a heterogenous reproductive disorder that is often associated with metabolic dysfunction, as well as comorbidities such as pregnancy complications. Although metabolic traits like hyperinsulinemia (*i*.*e*., elevated insulin without hypoglycemia) likely exacerbate the reproductive and metabolic features of PCOS, the precise impacts of specific metabolic traits on PCOS pathogenesis, symptom severity, and comorbidity incidence are not known. The aim of our study was to investigate the relationships between insulin levels, PCOS-like traits, and pregnancy complications by limiting endogenous insulin production in a mouse model of PCOS. Using *Ins1*-null mice with modulated *Ins2* gene dosage (*Ins1*^*-/-*^*:Ins2*^*+/-*^ versus *Ins1*^*-/-*^*:Ins2*^*+/+*^ littermates), we longitudinally assessed metabolic and reproductive phenotypes in PCOS-like mice generated via prenatal anti-Müllerian hormone (PAMH) exposure. We observed mild reproductive characteristics of PCOS in PAMH mice of both genotypes, including increased anogenital distances, delayed puberty, and disrupted estrous cycling, but did not detect robust PAMH-induced metabolic changes across six months. In the absence of PAMH-aggravated metabolic dysfunction or hyperinsulinemia—even in mice fed a high-fat, high-sucrose diet—reducing *Ins2* gene dosage did not notably change most measured traits. However, high-fat, high-sucrose-fed PAMH pregnant dams exhibited a diminished pregnancy-induced insulinogenic response and a trend for reduced β-cell mass compared to control mice, together with superior blood glucose homeostasis despite the physiological challenges of pregnancy. Therefore, while *Ins1*-null PAMH mice did not manifest pronounced PCOS-like metabolic features, prenatal AMH exposure can cause shifts in metabolic homeostasis during pregnancy.

## Introduction

Polycystic ovary syndrome (PCOS) is a heterogeneous reproductive endocrinopathy that affects up to 20% of reproductive-age women [1]. Currently, PCOS is most often diagnosed by the Rotterdam criteria, which require the presentation of at least two out of three key phenotypes: 1) ovulatory dysfunction and menstrual irregularity, 2) biochemical or clinical hyperandrogenism, and 3) polycystic ovarian morphology [2]. PCOS is associated with many comorbidities, including infertility and pregnancy complications [3–5]. Beyond these reproductive system symptoms, PCOS often features a metabolic component that can include hyperinsulinemia (*i*.*e*., elevated levels of circulating insulin without hypoglycemia), insulin resistance, obesity, and dyslipidemia [3,4]. Lean PCOS subgroups may have distinct reproductive traits and are less likely to exhibit metabolic features, but non-obese PCOS patients can still have some underlying metabolic dysfunction [6–9]. Overall, the extent to which metabolic features contribute towards the reproductive symptoms of PCOS remains to be determined [7,10].

It is likely that insulin resistance and/or hyperinsulinemia, present in approximately 60-95% of women with PCOS, compounds the reproductive and metabolic characteristics of PCOS [11,12]. Insulin acts as a co-gonadotroph in the ovaries, working synergistically with luteinizing hormone (LH) to increase theca cell production of androgens and upregulate theca cell LH receptors [13,14]. Furthermore, insulin decreases hepatic production of sex hormone binding globulin, allowing for greater action of free testosterone [15]. In PCOS pathogenesis, increased levels and actions of LH and androgens contribute towards abnormalities in folliculogenesis and dominant follicle selection, resulting in increased numbers of growing pre-antral follicles that lead to a polycystic ovarian morphology [16]. Elevated anti-Müllerian hormone (AMH), which is secreted by immature follicles, signals centrally alongside the altered levels of ovary-produced hormones to further exacerbate GnRH neuron hyperactivity and LH secretion, thus perpetuating further hyperandrogenism [17–19]. Since testosterone can itself increase insulin secretion and induce insulin resistance in certain tissues, the cycle of hyperinsulinemia, insulin resistance, and hyperandrogenemia is challenging to disentangle [6,20,21]. It has not yet been determined whether curtailing insulin secretion is sufficient to alleviate the severity of PCOS pathophysiology.

Metabolic features such as hyperinsulinemia are also implicated in some of the pregnancy complications experienced by PCOS patients. For instance, offspring of PCOS mothers have increased risk of preterm delivery, fetal growth restriction, and low birth weight [22]. Interestingly, experimentally inducing hyperinsulinemia in pregnant rats lowers maternal glucose levels and restricts fetal growth, highlighting the importance of maternal metabolic regulation for fine-tuned fetal nutrient delivery [23]. PCOS is associated with aberrations in maternal metabolism, such as gestational diabetes, which occurs in approximately 20% of PCOS mothers [24]. Gestational diabetes is defined as hyperglycemia, or elevated blood glucose levels, that develops during pregnancy and resolves following birth [25]. This occurs when there is inadequate β-cell compensation for the insulin resistance that occurs during pregnancy, leading to persistent hyperglycemia [26]. Notably, hyperinsulinemia prior to or during early pregnancy is a predictor of gestational diabetes [27,28]. While insulin sensitivity and insulin levels are important players in governing maternal and fetal health during pregnancy, questions remain about their contributions in the context of PCOS.

The aim of our study was to investigate the relationships between insulin levels, PCOS pathogenesis, and pregnancy complications by limiting endogenous insulin production in a PCOS-like mouse model [29]. We induced PCOS-like phenotypes via prenatal AMH (PAMH) treatment of mice with full or partial *Ins2* expression, on a background with complete ablation of the rodent-specific *Ins1* insulin gene (*Ins1*^*-/-*^*:Ins2*^*+/-*^ versus *Ins1*^*-/-*^*:Ins2*^*+/+*^ littermates). Unexpectedly, although we observed mild reproductive characteristics of PCOS in the PAMH mice of both genotypes, there were no overt PAMH-induced metabolic changes by six months of age, even in mice fed a high-fat, high-sucrose diet. In the absence of evident metabolic dysfunction or hyperinsulinemia, the effects of reducing *Ins2* gene dosage in *Ins1-*null PAMH mice were subtle. However, despite presenting with lean, non-metabolic PCOS-like phenotypes, the PAMH mice showed metabolic differences during pregnancy, including a suppression of the typical pregnancy-associated elevation in insulin levels and superior gestational glucose tolerance. Therefore, elevated prenatal AMH exposure may shift metabolic homeostasis during pregnancy, even in lean PCOS-like mice that did not have evident metabolic dysfunction previously.

## Results

### PAMH exposure did not induce severe metabolic dysfunction by 6 months of age in *Ins1-* null mice

To characterize the effects of insulin production on PCOS pathogenesis, we examined reproductive and metabolic traits of PCOS-like mice in littermates with full or partial *Ins2* gene dosage, on an *Ins1-*null background to prevent the compensatory *Ins1-*upregulation that can occur with reduced *Ins2* expression [30]. This strain was previously shown to generate female *Ins1*^*-/-*^*:Ins2*^*+/-*^ mice with lower fasting insulin than their *Ins1*^*-/-*^*:Ins2*^*+/+*^ littermates, a suppression of hyperinsulinemia that proved protective against high-fat diet-induce obesity and age-related metabolic decline [31,32]. Here, PCOS-like mice and control groups were generated with prenatal AMH or prenatal PBS (PPBS) exposure, via intraperitoneal injections of dams for three consecutive days late in the gestational period (Figure 1A). Prenatal AMH exposure was previously shown to induce hyperinsulinemia, hyperglycemia, impaired glucose tolerance, and reduced insulin sensitivity in PCOS-like female mice by 6 months of age [33]. Therefore, we hypothesized that reducing *Ins2* gene dosage would lower fasting insulin and improve metabolic health in PAMH mice.

**Figure 1.**
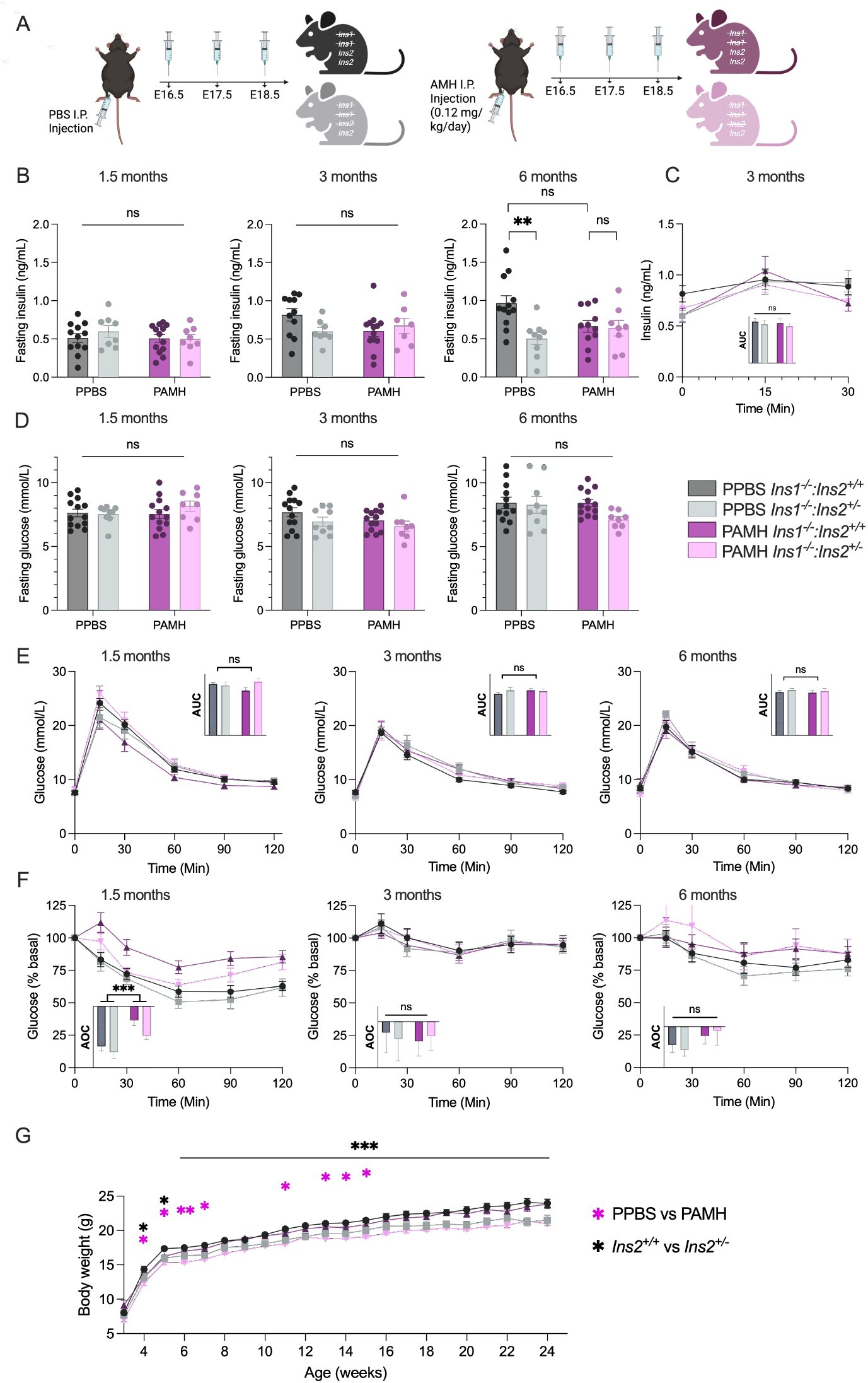
*Ins1-*null PAMH mice did not show overt metabolic changes or elevated insulin across 6 months of age, regardless of *Ins2* gene dosage. (A) Experimental mice were female *Ins1*^*-/-*^*:Ins2*^*+/+*^ and *Ins1*^*-/-*^*:Ins2*^*+/-*^ littermates that had been prenatally exposed to AMH (PAMH) or PBS (PPBS). *Ins1*^*-/-*^ *:Ins2*^*+/+*^ dams were given an intraperitoneal injection of 0.12 mg/kg/day AMH (n = 7 dams) or PBS (n = 7 dams) from E16.5 to E18.5 to generate PCOS-like mice and controls (PPBS *Ins1*^*-/-*^*;Ins2*^*+/+*^ n = 13; PPBS *Ins1*^*-/-*^*;Ins2*^*+/-*^ n = 9; PAMH *Ins1*^*-/-*^*;Ins2*^*+/+*^ n = 13; PAMH *Ins1*^*-/-*^*;Ins2*^*+/-*^ n = 8). (B) Periodic 4-h fasted plasma insulin levels (ng/mL) at three time points, as well as (C) measurements of glucose-stimulated insulin secretion over 30 minutes in 3 month-old mice. (D) Periodic 4-h fasted blood glucose levels (mmol/L) at three time points. Periodic measurements of blood glucose response to an intraperitoneal injection of (E) glucose (2g/kg) and (F) an insulin analog (0.75U/kg) over two hours, at 6-8 weeks, 3 months, and 6 months of age. Area under the curve (AUC; panel E; y-axis units of mmol/L• min) or area over the curve (AOC; panel F; y-axis units percent basal•min) is shown in panel insets. (G) Body mass measured across 6 months. Data are means ± SEM. Scatter points represent values from individual mice. Two-way ANOVA models were used to assess factors of genotype and AMH treatment, a significant interaction led to individual one-way ANOVAs with Bonferroni corrections. ns p>0.05, *p≤0.05, **p≤0.01, ***p≤0.001.

4h-fasted blood glucose and plasma insulin levels were intermittently measured across 6 months. No significant changes in insulin or glucose levels were observed in PAMH mice compared to PPBS controls during this time (Figure 1B, C, D). At 6 months of age, control PPBS *Ins1*^*-/-*^*:Ins2*^*+/-*^ mice had significantly reduced fasting insulin levels compared to their *Ins1*^*-/-*^ *:Ins2*^*+/+*^ littermates (p≤0.01; Figure 1B); however, no changes in fasting insulin were observed between PAMH *Ins1*^*-/-*^*:Ins2*^*+/+*^ and *Ins1*^*-/-*^*:Ins2*^*+/-*^ littermates. We also did not detect any differences in glucose-stimulated insulin secretion at 3 months due to PAMH exposure or *Ins2* gene dosage (Figure 1C). This indicates that neither fasting nor fed hyperinsulinemia was induced in *Ins1*-null PAMH mice. Unfortunately, without PCOS-like hyperinsulinemia we were unable to model a mitigation in elevated insulin production by reducing *Ins2* gene dosage in our experimental design.

Consistent with their lack of hyperinsulinemia, PAMH-exposed *Ins1-*null mice did not show other robust or persistent signs of metabolic dysfunction. Glucose tolerance tests revealed no differences between experimental groups in blood glucose responses to a glucose load at 1.5, 3, or 6 months (Figure 1E). Insulin tolerance tests revealed a transient reduction in insulin sensitivity at 1.5 months in PAMH mice (p≤0.001 for PAMH effect, Figure 1F). This was partially attenuated with reduced *Ins2* gene dosage, although the genotype effect did not reach the threshold for statistical significance (p = 0.0536 for *Ins1*^*-/-*^*:Ins2*^*+/-*^ versus *Ins1*^*-/-*^*:Ins2*^*+/+*^, Figure 1F). These metabolic differences were not sustained at 3 or 6 months of age, when there were similar blood glucose responses to insulin across all groups. Body mass was consistently reduced in both control and PAMH *Ins1*^*-/-*^*:Ins2*^*+/-*^ mice compared to *Ins1*^*-/-*^*:Ins2*^+/+^ littermates (p≤0.001; Figure 1G). There was also an early mild reduction of body mass in PAMH mice compared to controls from 3-15 weeks (p≤0.05; Figure 1G), though this effect may be due to litter-specific variations in body mass, which is common in C57BL/6J litters [34]. Overall, PAMH treatment in *Ins1-*null mice did not induce strong metabolic dysfunction beyond a transient reduction in insulin sensitivity during adolescence. Partial *Ins2* gene dosage may have mildly attenuated this brief period of insulin resistance and led to reduced weight gain, but it did not cause any notable changes in other measured parameters of metabolic health.

### Reproductive aberrations in PAMH mice were not improved by reduced *Ins2* gene dosage

PAMH exposure has been previously shown to induce all three key PCOS-like phenotypes in adult female mice (*i*.*e*., hyperandrogenism, ovulatory dysfunction, polycystic ovarian morphology) [29]. Since the precise contribution of insulin to the development and/or degree of these phenotypes has not been established, we assessed the effects of lowering *Ins2* gene dosage on various reproductive PCOS-like traits in PAMH mice.

PAMH exposure inhibits placental aromatase, leading to fetal hyperandrogenism that drives many of the neuroendocrine alterations that induce PCOS-like symptoms in adulthood [35]. To confirm fetal testosterone exposure, we assessed anogenital distance from postnatal day 25 to 60. PAMH mice had longer anogenital distances than PPBS controls (p≤0.001; Figure 2A). PAMH *Ins1*^*-/-*^*:Ins2*^*+/-*^ mice exhibited a trend for slightly smaller anogenital distances compared to full-*Ins2* littermates, suggesting that fetal hyperandrogenemia may be moderately diminished with reduced *Ins2* dosage; however, this did not reach the threshold for a statistically significant genotype effect (*e*.*g*., p = 0.0981 for *Ins1*^*-/-*^*:Ins2*^*+/-*^ versus *Ins1*^*-/-*^*:Ins2*^*+/+*^ at 35 days, Figure 2A). Both *Ins1*^*-/-*^*:Ins2*^*+/+*^ and *Ins1*^*-/-*^*:Ins2*^*+/-*^ PAMH mice displayed a slightly delayed time of vaginal opening compared to controls (p≤0.01; Figure 2B), indicating delayed pubertal onset [36].

**Figure 2.**
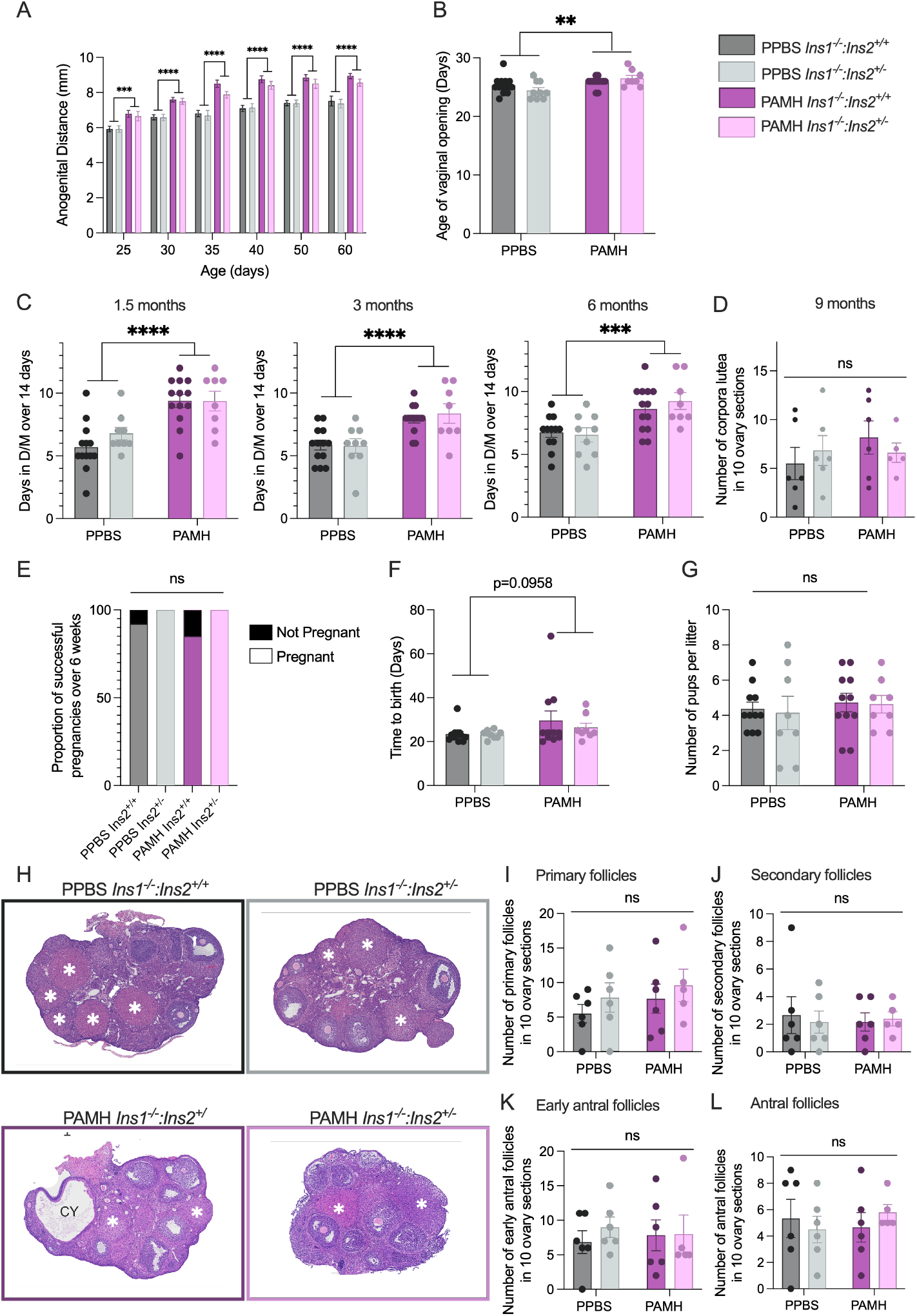
*Ins2* gene dosage did not significantly change PAMH-induced reproductive features. Anogenital distance measurements from postnatal day 25 to 60 in PAMH and PPBS mice with full or partial *Ins2* expression (PPBS *Ins1*^*-/-*^*;Ins2*^*+/+*^ n = 13; PPBS *Ins1*^*-/-*^*;Ins2*^*+/-*^ n = 9; PAMH *Ins1*^*-/-*^*;Ins2*^*+/+*^ n = 13; PAMH *Ins1*^*-/-*^*;Ins2*^*+/-*^ n = 8). (B) Time of vaginal opening, assessed daily from weaning. (C) Periodic analysis of proportion of time spent in metestrous and diestrous over 14 consecutive days, measured via analysis of vaginal cytology from 5-7 weeks, 10-12 weeks, and 23-25 weeks. (D) Numbers of corpora lutea across 10 serial ovarian sections per mouse. (E) Proportion of female mice that successfully became pregnant over 6-weeks of pairing with a wildtype male mouse. (F) Time-to-birth, measured from day of breeding pair setup with a wildtype 2-3-month-old male to day of birth. (G) Number of pups per litter. (H) Representative images of H&E-stained ovaries from 9-month-old PPBS and PAMH *Ins1*^*-/-*^*;Ins2*^*+/+*^ and *Ins1*^*-/-*^*;Ins2*^*+/-*^ mice (n = 6 per group). Asterix (*) indicate corpora lutea and CY indicates a cyst. (I) Number of primary follicles per ovary. (J) Number of secondary across 10 serial ovarian sections per mouse. (K) Number of early antral follicles across 10 serial ovarian sections per mouse. (L) Number of antral follicles across 10 serial ovarian sections per mouse. Corpora lutea and follicles were counted over 10 serial sections taken 50μm apart per ovary. Data are means ± SEM. Scatter points represent values from individual mice. Two-way ANOVA models were used to assess factors of genotype and AMH treatment, and a Fisher’s exact test was used to assess the proportion of successful pregnancies. ns p>0.05, *p≤0.05, **p≤0.01, ***p≤0.001, ****p≤0.0001.

To assess ovulatory function, we examined estrous cycling and corpora lutea (CL). PAMH mice exhibited dysregulation of their estrous cycles compared to controls, consistently spending more time in metestrous and diestrous (*i*.*e*., the non-ovulatory phases of the estrous cycle) over a 14-day period compared to PPBS controls (p≤0.001; Figure 2C). This effect was not attenuated by partial *Ins2* gene dosage. We did not observe complete anovulation in PAMH mice, as they did enter the ovulatory phases of the estrous cycle and had similar CL numbers to PPBS controls at 9 months of age (Figure 2D). PAMH mice also did not have significant fertility disruptions, reflected by similarities in the proportion of successful pregnancies, times to birth, and number of pups per litter, compared to PPBS controls (Figure 2E, F, G). PAMH mice tended to have slightly increased latency to birth after being paired with a male, though this was not a statistically significant effect (p = 0.0958; Figure 2F). Further analysis of ovarian morphology following breeding trials revealed no significant changes in numbers of primary, secondary, early antral, or antral follicles in PAMH mice (Figure 2H, I, J, K, L). Large cysts with thin granulosa cell layers and large antral follicles (*e*.*g*. Figure 2H) were observed in ovaries of both PAMH and control mice, though the presence of these cysts was infrequent across all groups.

Taken together, these data demonstrate that *Ins1-*null PAMH mice displayed mild reproductive PCOS-like phenotypes, including elongated anogenital distances, delayed time of vaginal opening, and disrupted estrous cycles. However, PAMH exposure did not significantly alter fertility or ovarian morphology. Reducing *Ins2* gene dosage did not have appreciable effects on timing of puberty or estrous cycling, though a trend for PAMH *Ins1*^*-/-*^*:Ins2*^*+/-*^ mice to have slightly smaller anogenital distances than *Ins1*^*-/-*^*:Ins2*^*+/+*^ littermates indicates the possibility that *Ins2* might heighten fetal hyperandrogenism.

### Gestational glucose homeostasis was altered in high-fat, high-sucrose-fed PAMH mice

Given the associations between PCOS and such pregnancy complications as gestational diabetes and fetal growth restriction, we wished to assess how prenatal AMH treatment affects maternal and fetal health metrics during the metabolically demanding state of pregnancy. Due to the absence of strong metabolic aberrations in PAMH mice on a chow diet (Figure 1), mice were first exposed to a high-fat, high-sucrose (HFHS) diet beginning at postnatal day 75 (∼2.5 months of age), with a goal of generating an obese PCOS-like phenotype. Following one month of HFHS diet, fasting blood glucose and plasma insulin levels, and glucose-stimulated insulin secretion were not altered by PAMH exposure or *Ins2* gene dosage (Figure 3A, B, C). This was consistent with observations from 3-month-old chow-fed mice (Figure 1). After 2.5 months of HFHS diet, fasting glucose levels were significantly reduced in *Ins1*^*-/-*^*:Ins2*^*+/-*^ PAMH and PPBS mice compared to littermates with full *Ins2* expression (p < 0.05; Figure 3D), although there were no significant differences in fasting insulin levels (Figure 3E). This indicates that PAMH exposure did not worsen hyperinsulinemia or hyperglycemia in HFHS-fed mice, relative to HFHS-fed PPBS controls. At 5 months of age, HFHS-fed PAMH mice had irregular estrous cycles compared to PPBS mice (p≤0.01; Figure 3F), consistent with the effects of PAMH exposure on a chow diet (Figure 2). A glucose tolerance test revealed that all four HFHS-fed groups had sustained glucose peaks that were delayed to 30 minutes post-glucose load, suggestive of some degree of glucose intolerance that was not exacerbated by PAMH exposure or attenuated by reduced *Ins2* gene dosage (Figure 3G). Similarly, insulin sensitivity was not significantly altered by PAMH exposure or *Ins2* gene dosage (Figure 3H). Body mass was lower in *Ins1*^*-/-*^*:Ins2*^*+/-*^ mice compared to *Ins1*^*-/-*^: *Ins2*^*+/+*^ littermates for both PAMH and PPBS treatments; there was no effect of PAMH exposure on body weight over 6 months (Figure 3I). These data suggest that 2.5 months of HFHS diet induced some aberrations in metabolic health, but this effect was not exacerbated by PAMH exposure. There was a mild protective effect of lowering *Ins2* gene dosage on fasting glucose levels and body mass; however, no notable alterations in insulin levels or insulin sensitivity were observed alongside these effects.

**Figure 3.**
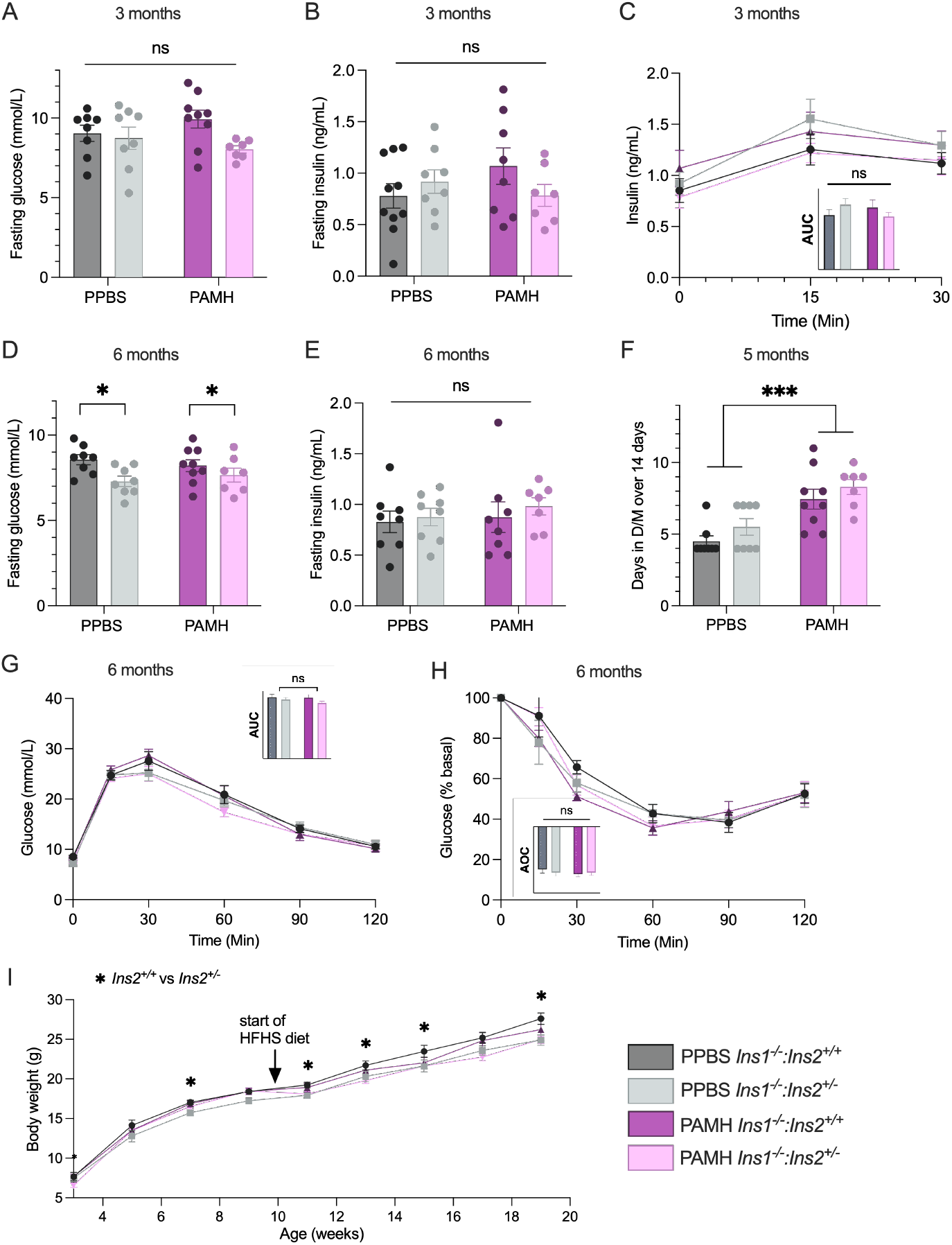
Metabolic dysfunction not exacerbated by prenatal AMH exposure in mice on a high-fat, high-sucrose diet. (A) 4-h fasted blood glucose levels (mmol/L) and (B) 4h-fasted plasma insulin levels (ng/mL) at 3 months of age, in HFHS diet-fed PAMH and PPBS mice with partial or full *Ins2* gene dosage (PPBS *Ins1*^*-/-*^*;Ins2*^*+/+*^ n = 8; PPBS *Ins1*^*-/-*^*;Ins2*^*+/-*^ n = 8; PAMH *Ins1*^*-/-*^*;Ins2*^*+/+*^ n = 9; PAMH *Ins1*^*-/-*^ *;Ins2*^*+/-*^ n = 7). (C) Measurements of glucose-stimulated insulin secretion over 30 minutes at 15 weeks, with area under the curve (y-axis units ng/mL insulin•min) in panel inset. Data are means ± SEM. (D) 4-h fasted blood glucose levels (mmol/L) and (E) 4h-fasted plasma insulin levels (ng/mL) at 6 months of age. (F) Analysis of time spent in metestrous and diestrous over 14 consecutive days, measured from 18-20 weeks of age. (G) Blood glucose responses to an intraperitoneal injection of glucose (2g/kg) and (H) an insulin analog (1.0 U/kg) in female PAMH and PPBs mice at 6 months of age. Area under the curve (AUC; panel G; y-axis units of mmol/L•min) or area over the curve (AOC; panel H; y-axis units percent basal•min) is shown in panel insets. (I) Body mass (g) measured over 6 months. Data are means ± SEM. Scatter points represent values from individual mice. Two-way ANOVA models were used to assess factors of genotype and AMH treatment. ns p>0.05, *p≤0.05, **p≤0.01, ***p≤0.001, ****p≤0.0001.

Interestingly, despite the absence of overt PAMH-induced metabolic phenotypes in our mice—even on a high-fat, high-sucrose diet—we found that metabolic differences between PAMH and PPBS groups did become evident during pregnancy. PAMH *Ins1*^*-/-*^: *Ins2*^*+/+*^ mice tended to have the lowest proportion of successful pregnancies of these HFHS-fed groups (Figure 4A), paralleling the results of the breeding trial of the chow-fed mice (Figure 2E), but this was not statistically significant (p = 0.2941 for PAMH *Ins1*^*-/-*^: *Ins2*^*+/+*^ vs. PPBS *Ins1*^*-/-*^: *Ins2*^*+/+*^ groups, Figure 4A). Of the mice that did become pregnant, we observed that only control PPBS mice showed the expected elevations in mean fasting insulin during pregnancy (at G14.5), compared to pre-pregnancy levels (p<0.01; Figure 4B), whereas PAMH mice did not display a significant pregnancy-induced rise in insulin levels (p = 0.1278; Figure 4B). Gestational diabetes often occurs due to inadequate β-cell compensation for the insulin resistance that develops during pregnancy, resulting in maternal hyperglycemia and insulin insufficiency [26]. Examination of β-cell mass at G14.5 suggested a trend for reduced β-cell mass of PAMH mice compared to PPBS controls (p = 0.0660; Figure 4C, D); this was not affected by *Ins2* gene dosage. This could have contributed towards the suppressed pregnancy-induced escalation in insulin levels observed in the PAMH mice. Unexpectedly, however, PAMH mice had significantly superior glucose tolerance at G14.5 compared to PPBS controls (p≤0.001; Figure 4E); again, there was no genotype effect. These maternal metabolic changes did not appear to alter measured parameters of fetal health at G14.5, demonstrated by consistent number of fetuses per dam (Figure 4F), fetal-to-placental weight ratios (Figure 4G), and fetal weights (Figure 4H) across groups.

**Figure 4.**
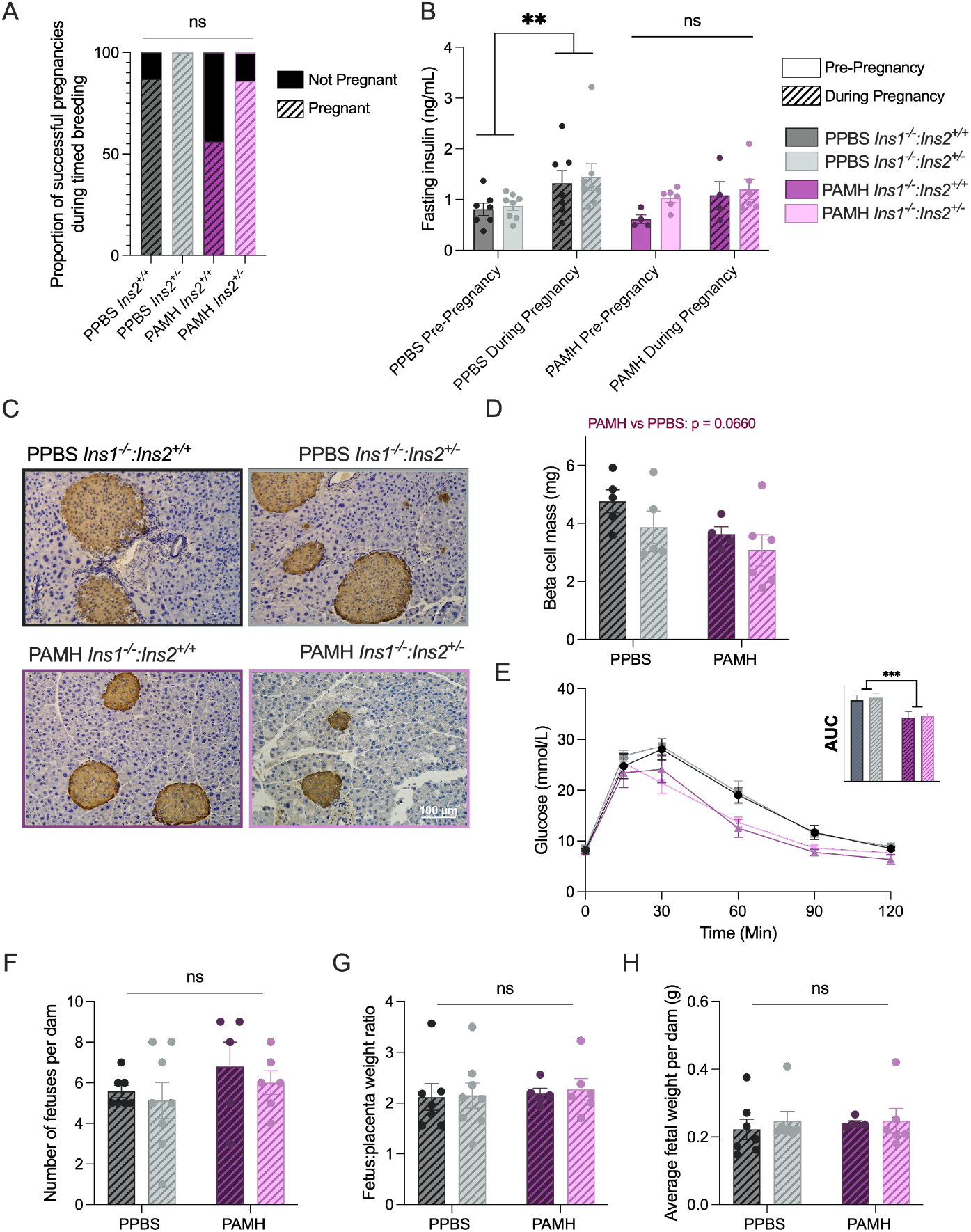
Pregnant PAMH mice showed suppression of pregnancy-induced insulin elevation and superior gestational glucose tolerance. (A) Proportion of HFHS-fed female mice that successfully became pregnant in timed breeding trials with a wildtype male mouse (PPBS *Ins1*^*-/-*^*;Ins2*^*+/+*^ n = 8; PPBS *Ins1*^*-/-*^*;Ins2*^*+/-*^ n = 8; PAMH *Ins1*^*-/-*^*;Ins2*^*+/+*^ n = 9; PAMH *Ins1*^*-/-*^*;Ins2*^*+/-*^ n = 7). (B) 4-h fasting plasma insulin levels (ng/mL) pre-pregnancy in those 6-month-old mice that successfully conceived in the timed breeding trials, compared to the 4-h fasting plasma insulin levels at G14.5 of pregnancy (PPBS *Ins2*^*+/+*^ n=7; PPBS *Ins2*^*+/-*^ n=8; PAMH *Ins2*^*+/+*^ n=4; PAMH *Ins2*^*+/-*^ n=6; pre-pregnancy fasting insulin levels are a subset of those shown in Figure 3E). (C) Representative images of DAB (3’3-diaminobenzidine) staining for insulin (brown) with hematoxylin counterstaining (blue) in pancreata of PAMH and PPBS mice at G14.5. Images were taken at 20x magnification using Brightfield microscopy. (D) Quantification of β-cell mass, calculated by multiplying %β-cell area by mass of pancreas. n=4-6 per group, five pancreatic sections were quantified per animal. (E) Blood glucose response to an intraperitoneal injection of glucose (2g/kg) at G14.5. Area over the curve (y-axis units of mmol/L•min) is shown in panel inset. (F) Number of fetuses per dam. (G) Fetal:placental mass ratio at G14.5. Calculated from average fetus:placenta mass ratios of each dam. (H) Average fetal weight per dam (g), calculated from average weight of litter. Two-way ANOVA models were used to assess factors of genotype and AMH treatment, and a Fisher’s exact test was used to assess the proportion of successful pregnancies. Repeated measures ANOVAs were used to compare pre-pregnancy and mid-pregnancy insulin levels for PPBS and PAMH mice; the effects of *Ins2* genotype were not statistically significant in either case. Mixed model analysis (with dam as a random factor) were used to assess fetal traits where there were multiple offspring per dam. ns p>0.05, *p≤ 0.05, ***p≤0.001.

These data suggest that PAMH mice may have had reduced β-cell hyperplasia and/or hypertrophy during pregnancy, resulting in suppression of compensatory insulin production. Interestingly, this was not associated with hyperglycemia or abnormal glucose homeostasis during a glucose tolerance test at G14.5. In fact, PAMH mice showed superior blood glucose disposal in response to a glucose load compared to PPBS controls. None of these effects were significantly altered by reduction of *Ins2* gene dosage.

## Discussion

In this study, we induced PCOS-like symptoms via prenatal-AMH exposure in *Ins1*-null mice with full or partial *Ins2* expression. We found that while *Ins1-*null PAMH mice displayed some reproductive characteristics consistent with a lean PCOS model, they did not show robust phenotypes of metabolic dysfunction over the course of six months. The reproductive phenotypes we observed also tended to be less pronounced than other reported examinations of the PAMH mouse model [33,35]. Phenotypes of animal models can also be affected by many genetic and/or environmental factors, such as animal facility or housing effects, mouse strain, and differences in peptide reagent quality or preparation (*e*.*g*. slight differences in lyophilized AMH or its reconstitution) [37]. Considering that we did not detect robust metabolic defects in PAMH mice, it is also possible that metabolic dysfunction might have further augmented the reproductive features of PCOS.

In the absence of evident PCOS-associated features of metabolic dysfunction like hyperinsulinemia, we could not effectively test or observe outcomes that might have resulted from suppressing insulin hypersecretion. *Ins1*-null PAMH mice transiently exhibited reduced insulin sensitivity at 8 weeks of age, potential due in part to PAMH-induced fetal androgenization [38,39], but hyperinsulinemia was not successfully induced in this PCOS-like model (compared to PPBS controls). Moreover, while a metabolically unhealthy HFHS diet generated relatively high fasting insulin and glucose intolerance for a separate cohort of PAMH and PPBS mice, this response was not exacerbated by prenatal AMH exposure. Our results show discrepancy from other prenatal androgenization mouse models of PCOS, which often generate noticeable metabolic dysfunction, including hyperinsulinemia and insulin resistance [40,41]. Indeed, previous work with PAMH mice revealed 2-fold higher fasting insulin levels, a ∼30% increase in fasting glucose levels, greater glucose-stimulated insulin secretion, and reduced glucose tolerance and insulin sensitivity by 6 months of age [33,42].

Notably, our experimental animals were all on an *Ins1*-null background, and this may have affected the generation of PCOS-like metabolic phenotypes. While humans have a single insulin gene, mice possess two non-allelic insulin genes—rodent-specific *Ins1*, and the ancestral insulin gene, *Ins2* [43]. Although the *Ins1* and *Ins2* genes are incompletely characterized, there are differences in tissue-specific and temporal expression patterns which suggest that *Ins1* and *Ins2* may be differentially regulated [44]. There also appear to be sex-dependent phenotypes that arise from manipulating these two genes. For example, while reducing *Ins1* dosage effectively lowered circulating insulin levels in *Ins2*-null male mice (compared to *Ins*^*+/+*^*:Ins2*^*-/-*^ male littermates that developed high-fat diet-induced hyperinsulinemia with full *Ins1* expression), the female *Ins2*-null littermates failed to develop high-fat diet-induced hyperinsulinemia regardless of *Ins1* gene dosage [45]. Conversely, female but not male *Ins1*^*-/-*^*:Ins2*^*+/-*^ mice showed significantly attenuated high-fat diet-induced hyperinsulinemia due to reduced *Ins2* dosage [32,46]. This suggests that *Ins1* expression may be a more sensitive and tuneable contributor to hyperinsulinemia in male mice, which could potentially be driven by androgen hormones if they have disparate regulatory effects on *Ins1* versus *Ins2*. PAMH exposure is believed to induce a fetal hyperandrogenic environment, which causes neuroendocrine reprogramming in rodent female pups leading to hyperandrogenism and other defects in adulthood [35]. Mishra *et al*., (2018) showed that exposure to dihydrotestosterone (DHT) significantly increased *Ins1* and *Ins2* mRNA expression in cultured female rat islets, but it was noted that *Ins1* may have been the greater contributor to the rise in insulin secretion; *Ins1* mRNA was greater than doubled in response to DHT, while *Ins2* mRNA expression only exhibited a 0.25-fold increase [20].

Similarly, Benrick *et al*. (2017) observed that DHT increased the expression of *Ins1* mRNA but not *Ins2* mRNA in the pancreata of adiponectin knockout mice [47]. Therefore, while more work is needed to test this conjecture, it is possible that high levels of androgens might impart some of their insulinogenic effect in rodents through preferential stimulation of *Ins1* expression compared to *Ins2*. This could occur via a functional androgen response element that has been identified in the *Ins1* promoter region [20].

Pregnancy requires extreme endocrine, metabolic, and physiological changes that can expose underlying features of metabolic dysfunction, so we wished to evaluate whether the challenges of pregnancy might reveal masked metabolic features in these *Ins1*-null PAMH mice. During pregnancy, placenta-produced hormones cause maternal insulin resistance and hepatic glucose production to ensure adequate glucose supply for fetal growth during late pregnancy stages. To maintain sufficient maternal nutrient stores despite this fetal demand, these effects are counterbalanced maternally by an expanded β-cell pool and increased insulin production [48]. In rodents, maternal β-cell adaptation during gestation is not generated as a reactive response to pregnancy-induced insulin resistance, but instead seems to precede insulin resistance in anticipation of these increased maternal nutrient requirements [49,50]. Many of these critical metabolic responses to pregnancy are influenced by a suite of hormones [48]. Levels of estradiol, progesterone, cortisol, and prolactin increase drastically during pregnancy, and have been shown to cause post-binding defects in insulin action [51,52]. For instance, mice with progesterone receptor-deficiency demonstrate increased insulin sensitivity during pregnancy [53]. Considering that PAMH mice have been shown to have a baseline reduction in progesterone levels compared to controls [35,42], it seemed possible that prenatal AMH exposure could perpetuate long-lasting endocrine disturbances that might shift metabolic function during pregnancy.

Indeed, we found that prenatal AMH exposure culminated in irregular responses to pregnancy for the HFHS-fed mice in our study. Pregnant PAMH dams tended to show a reduction in gestational β-cell mass compared to PPBS controls. Likewise, the expected pregnancy-associated rise in circulating insulin that appeared by G14.5 in PPBS mice was not evident in PAMH mice. This suggests that prenatal AMH may have led to impairment of β-cell adaptation during pregnancy in HFHS-fed mice. This did not correspond with glucose intolerance, which would be indicative of gestational diabetes. Instead, pregnant PAMH mice had superior glucose tolerance compared to PPBS controls at G14.5, which might have been linked to a preservation of maternal insulin sensitivity despite pregnancy. Therefore, while PAMH mice did not present with a gestational diabetes phenotype in our study, we postulate that prenatal AMH exposure may induce irregularities in gestational metabolism that might have eventually altered fetal nutrient supply and/or maternal nutrient stores during late gestation.

In this study, we observed that *Ins1-*null PAMH mice showed some lean PCOS-like reproductive features without pronounced metabolic dysfunction. In the absence of metabolic defects, particularly hyperinsulinemia, reducing *Ins2* gene dosage did not exert noteworthy effects on measured metabolic or reproductive characteristics aside from body mass, although slight trends suggested subtle effects of *Ins2* on early insulin sensitivity and anogenital distance. However, closer examination of pregnancy in six-month-old, high-fat, high-sucrose-fed mice revealed that PAMH mice had a trend for reduced β-cell mass and diminished pregnancy-induced elevation of insulin compared to PPBS mice, together with superior blood glucose homeostasis that points to preservation of their insulin sensitivity. This differs from expected maternal metabolic responses to pregnancy. Thus, despite the subtleties of PCOS-like metabolic features in *Ins1-*null PAMH mice, prenatal AMH exposure can cause shifts in metabolic homeostasis during pregnancy, irrespective of *Ins2* gene dosage. These findings highlight the potential for a lean, non-metabolic clinical subtype of PCOS to exhibit underlying metabolic features under the demands of pregnancy.

## Materials and Methods

### Experimental Animals

All animal procedures were approved by the University of Victoria Animal Care Committee (AUP#2021-017), following Canadian Council for Animal Care Guidelines. Experiments were carried out using *Ins1-*null female mice with genetically reduced *Ins2* gene dosage (*Ins1*^*-/-*^*:Ins2*^*+/-*^) and corresponding female littermate controls with full *Ins2* expression (*Ins1*^*-/-*^*:Ins2*^*+/+*^). *Ins1*^*-/-*^*:Ins2*^*+/-*^ and *Ins1*^*-/-*^*:Ins2*^*+/+*^ mice were previously established on a mixed background that is primarily (>90%) C57BL/6J (strain provided to our lab by J. Johnson), and this mouse colony was maintained by interbreeding *Ins1*^*-/-*^*:Ins2*^*+/+*^ and *Ins1*^*-/-*^*:Ins2*^*+/-*^ mice. All breeding trials or timed pregnancies with experimental animals used wild-type C57BL/6J males (strain #000664, The Jackson Laboratory, ME, USA). Mice were fed a chow diet (PicoLab Rodent Diet 20; 20% protein, 4% fat) from weaning at P21, and a subset of mice were assigned a high-fat, high-sucrose (HFHS) diet at P75 (45 kcal% fat; Research Diets D12451 irradiated; Research Diets, NB). All animals were housed in the University of Victoria Animal Care Facility at ∼21°C, on a 12:12hr light:dark cycle. Mice were group-housed for most experiments, except when separated to single housing for the duration of breeding and pregnancy.

### Generation of PCOS-like phenotypes

PCOS-like phenotypes were generated via prenatal exposure to anti-Müllerian hormone (AMH), following methods outlined in Mimouni & Giacobini [29]. Timed breeding pairs were established by assigning 2-4 month old *Ins1*^*-/-*^*:Ins2*^*+/-*^ males to singly housed 3-4 month old *Ins1*^*-/-*^*:Ins2*^*+/+*^ females. One male was placed in the cage of each female approximately 1 hour before the animal facility lights were turned off. The following morning, the presence of a vaginal plug was assessed between 7:00 and 9:00am, and the male was removed from the cage. If a vaginal plug was not observed, this process was repeated over four days until all females displayed a vaginal plug. The day of vaginal plug observation was assigned embryonic day 0.5 (E0.5). Pregnancy was confirmed by a >10% weight increase one week following copulation, and by visual assessment and palpation on E14.5.

Lyophilized AMH (Recombinant Human MIS/AMH Protein, Bio-Techne, MN, USA, Cat #1737-MS-010) was reconstituted in sterile reconstitution buffer (4mM HCl diluted in phosphate buffered saline (1X PBS; pH 7.4; Gibco, Cat #10010-023) containing 0.1% bovine serum albumin (VWR, Cat #0332-25G)). From E16.5 to E18.5 dams were given an intraperitoneal (i.p.) injection of 200μL of freshly prepared AMH (0.12 mg/kg/day) between 8:00 and 9:00 am to generate the prenatal-AMH (PAMH) group, or with 200μL reconstitution buffer diluted in PBS to generate the control (PPBS) group (Figure 1A). Offspring were weaned at postnatal day 21 (P21) onto a normal chow diet. PAMH and PPBS mice were generated over separate cohorts. The set of mice were used to examine PCOS pathophysiology, while a separate set was fed a HFHS diet and used to examine health during pregnancy.

#### *In vivo* reproductive phenotypes

Following weaning, PAMH and PPBS mice were observed daily for vaginal opening via visual assessment of the vulva. To confirm fetal testosterone exposure, anogenital distance was measured from the top of the anus to the bottom of the vagina, beginning at P25 and continuing throughout adolescence (P30, 35, 40, 50, 60).

To assess estrous cyclicity, vaginal smears were collected daily between 8:00 and 9:00am for 14 consecutive days. Cells were stained with 0.1% crystal violet (ThermoFisher Scientific, Cat #B12932.14; dissolved in ddH_2_O). Estrous cycle stage was determined by the presence and density of cornified squamous epithelial cells, leukocytes, and/or nucleated epithelial cells. Proestrous was identified by the presence of primarily nucleated epithelial cells, estrous was identified by the presence of mostly cornified squamous epithelial cells, metestrous was identified by the presence of all three cell types, and diestrous was identified by the presence of leukocytes and nucleated epithelial cells. Estrous cycling was performed for the first set of assessed mice at 5-7 weeks, 12-14 weeks, and 23-25 weeks. For HFHS-fed mice, estrous cycling was performed once, from 18-20 weeks of age.

To assess fertility and breeding capacity, a breeding trial was performed when mice were approximately six months old. Female mice were individually housed, and a single 2-3-month-old wild-type C57BL/6J male was added to each cage. Males remained in cages for six weeks, or until pregnancy was confirmed and maintained over a week.

### Ovarian histology

At nine months of age, chow diet-fed mice were euthanized via isoflurane inhalation and cervical dislocation. The left ovary was collected and fixed for 24 hours at 4°C in 4% paraformaldehyde (PFA, Electron Microscope Sciences, Cat#157-8). Standard paraffin embedding, sectioning, and hematoxylin and eosin (H&E) staining were conducted by the Molecular and Cellular Immunology Core at the Deeley Research Centre, BC Cancer, Victoria, BC. Ten 5μm sections were taken 50μm apart for each ovary. Each section was imaged using Brightfield microscopy at 40x magnification.

Numbers of corpora lutea as well as primary, secondary, early antral, and antral follicles were assessed in each ovary section by an investigator that was blinded to experimental group. Primary follicles were identified by the presence of a single layer of cuboidal granulosa cells. Secondary follicles were surrounded by greater than a single layer of cuboidal granulosa cells, with no observable antrum. To avoid repeat counting, only primary and secondary follicles that had a visible nucleolus in the section were counted. Antral follicles were counted in sections where there was a visible antral cavity, to avoid undercounting when nuclei were not imaged. To preclude repeat counting, the positions of early antral and antral follicles were marked and any follicles in the same position in subsequent sections were not counted. Early antral follicles were identified by emerging antral spaces, while antral follicles had a clearly defined antrum. Corpora lutea (CL) were identified by the presence of large, dense clusters of granulosa lutein cells with pale cytoplasm and smaller, more deeply stained theca lutein cells. CL are generally large and encompass multiple serial sections—to prevent repeat counting, CL were annotated, and their orientation and position was denoted. Any CL that had the same orientation and position in subsequent sections was not counted.

#### *In vivo* metabolic phenotypes

Mice were fasted for 4 hours in clean cages during the light period prior to any measurements of circulating factors. Blood glucose was measured using a OneTouch Ultra2 glucose meter (LifeScan Canada Ltd., Malvern, PA, USA). For intraperitoneal glucose tolerance tests (IPGTT) and intraperitoneal insulin tolerance tests (IPITT), blood glucose responses to an i.p. injection of glucose (2g/kg) or insulin analog (Humalog; 0.75U/kg for chow diet-fed mice at all time points, and 1.0 U/kg for HFHS-fed mice in 6 month-old IPITT) was measured over 2 hours. Intraperitoneal glucose-stimulated insulin secretion was measured over 30 minutes. Fasting plasma insulin and glucose-stimulated insulin levels were measured using a commercial mouse insulin ELISA kit (Alpco Diagnostics, Salem, NH, USA, Cat #80-INSMSU-E01) according to manufacturer protocols. Fasting glucose and insulin levels as well as blood glucose responses to an intraperitoneal delivery of glucose or insulin were measured three times over 6 months. Glucose-stimulated insulin secretion was measured at 14-weeks of age. In HFHS-fed mice, glucose-stimulated insulin secretion was measured at 15-weeks of age, followed by assessment of fasting glucose and insulin levels, IPGTT, and IPITT at 22 and 23 weeks, respectively.

### Gestational diabetes and fetal health

To exacerbate metabolic dysfunction, a subset of PAMH and PPBS mice were assigned to a HFHS diet at postnatal day 75, in adulthood. Metabolic health and reproductive characteristics were assessed in these mice as outlined above.

At 24 weeks of age, the HFHS-fed mice were paired with wild-type males for timed breeding. All mice were maintained on a HFHS diet during breeding and pregnancy. On G14.5, pregnant mice were fasted in clean cages for 4 hours during the light cycle. Fasting glucose and insulin were measured, and an IPGTT was performed. Following the IPGTT, pregnant mice were euthanized via isoflurane inhalation and cervical dislocation. The pancreas from each mouse was collected, weighed and immediately placed in cold 4% PFA. Each fetus and corresponding placenta were counted and weighed. Pancreata were fixed for 24 hours at 4°C in 4% PFA. Following fixation, standard tissue dehydration, paraffin embedding, and sectioning were conducted by the Molecular and Cellular Immunology Core at the Deeley Research Centre, BC Cancer, Victoria, BC. Six 5μm sections were taken 200μm apart for each pancreas.

### Insulin immunohistochemistry and β-cell mass measurement

Prior to immunohistochemistry, pancreas sections were deparaffinized and rehydrated. Heat-induced epitope retrieval was performed by microwaving tissues in sodium citrate buffer (10mM, 0.05% Tween 20, pH 6.0) for 10 minutes. Sections were incubated with 3% H_2_O_2_ for 10 minutes to quench endogenous peroxidase activity, this was immediately followed by blocking with normal goat serum (VECTASTAIN ABC Kit, Peroxidase (Guinea Pig IgG), Vector Laboratories Inc., Cat #PK-4007) for 20 minutes. Sections were incubated with insulin primary antibody (1:5; FLEX Polyclonal Guinea Pig Anti-Insulin, Agilent/Dako, Cat #IR002) overnight at 4°C.

Tissues were washed and incubated with biotinylated secondary antibody (Goat Anti-Guinea Pig IgG, biotinylated, part of the VECTASTAIN ABC Kit, Peroxidase (Guinea Pig IgG), Vector Laboratories Inc., Cat #PK-4007) for 1 hour in the dark at room temperature. Sections were incubated for 30 minutes in the dark with ABC reagent (VECTASTAIN ABC Kit, Peroxidase (Guinea Pig IgG), Vector Laboratories Inc., Cat #PK-4007). 1X peroxidase substrate solution (ThermoFisher Scientific, Cat #34002) was applied to sections for 10 minutes. Sections were stained with modified Mayer’s hematoxylin (Epredia, Cat #72804) and briefly immersed in bluing solution (0.003% NH_4_OH) to counterstain for nuclei. Sections were dehydrated and cleared in sequential ethanol and xylene washes, followed by cover-slipping with VectaMount Express Mounting Medium (Vector Laboratories Inc., Cat #H-5700-60).

Five sections per pancreas were imaged at 4x magnification using Brightfield microscopy. β-cell area was measured using ImageJ Colour Deconvolution 2 Software. Beta cell mass was calculated by multiplying the wet mass of the pancreas by the percent area of β-cells in each section.

### Imaging and Statistical Analyses

All image capturing was performed using Brightfield microscopy on a Nikon Eclipse Ti2 microscope. After image capture, all images were blinded and randomized prior to analysis. All statistical analyses and graphs were performed and created using GraphPad Prism 10 software. For most statistical analyses, two-way ANOVA or repeated ANOVA models were used to assess factors of PAMH exposure and genotype. A significant statistical interaction led to a one-way ANOVA comparison of groups with Bonferroni correction. For analyses of categorical variables (*i*.*e*., proportion of successful pregnancies), individual Fisher’s exact tests were used to compare incidences of the phenotype within experimental groups. Mixed model analyses (with dam as a random factor) were used to assess fetal traits where there were multiple offspring per dam. In all cases, the significance level was set at *p* ≤ 0.05.

